# Methionine γ-lyase Traps Cofactor Pyridoxal-5′-phosphate through Specific Serine-Mediated Affinity

**DOI:** 10.1101/2025.07.27.667015

**Authors:** Chang Yan, Xueting Liu, Ruiqi Su, Mengwen Shi, Zhi Geng, Ying Chen, Liwen Zhu, Zongjie Dai, Junfeng Hui, Xi Chen, Kai-Zhi Jia

## Abstract

Pyridoxal-5′-phosphate (PLP), the ubiquitous and ancient cofactor, plays important roles in enzymatic elimination, transamination and other reactions. The catalytic efficiency of PLP-dependent enzymes is significantly higher than that of free PLP. The recruitment of PLP from the environment by the enzymes, particularly through interactions outside the active site, is the key step determining the occurrence of PLP-mediated catalysis. However, the precise mechanism by which enzymes recruit PLP remains elusive. Methionine γ-lyase (MGL), a PLP-dependent enzyme, catalyzes the degradation of L-methionine, thereby suppressing cancer cell proliferation through serum or dietary methionine depletion. Here, we report the crystal structure of yMGL, which belongs to a newly identified subgroup of cystathionine γ-lyases, in complex with L-methionine and PLP. Through truncating the C-terminal domain of yMGL both *in vitro* and *in vivo*, we demonstrated that this domain, outside the canonical PLP-binding domain, is essential for the specific interaction between yMGL and PLP, as well as for efficient L-methionine catabolism. A conserved Ser in the C-terminal domain, positioned at the active-site entrance of MGLs, confers specificity for PLP. These findings elucidate a previously uncharacterized mechanism of PLP recruitment by MGLs and promote rational design of MGLs with tailored cofactor selectivity and catalytic performance.

## Introduction

Pyridoxal-5′-phosphate (PLP), the biologically active form of vitamin B6, serves as a ubiquitous and highly versatile cofactor in enzymatic catalysis, particularly in the metabolism of amino acids; the biosynthesis of antibiotics such as indolmycin and azomycin; and the production of key neurotransmitters, including serotonin, dopamine, epinephrine, norepinephrine, gamma-aminobutyric acid (GABA), and histamine.^[1–4]^ PLP-dependent enzymes catalyze elimination, transamination, decarboxylation, racemization, substitution, and retro-aldol cleavage reactions and are estimated to account for ∼ 4% of all enzyme activities.^[5]^ Despite extensive studies, the functional repertoire of PLP-dependent enzymes continues to expand, with newly discovered reactions and previously unannotated proteins being reported in recent years.^[6,7]^ The remarkable catalytic versatility of PLP arises from its unique chemical features. By acting as an “electron sink”, PLP delocalizes electron density through its conjugated pyridine ring via resonance, thereby stabilizing high-energy intermediates during catalysis.^[7]^ Central to its function is the formation of a Schiff base (internal aldimine) with a conserved lysine at the active site, which is subsequently displaced by the substrate to form an extermal aldimine.^[8]^ This transformation enables a variety of key reactions, including decarboxylation, deprotonation and carbanion stabilization through the quinonoid intermediate, to proceed under physiologically mild conditions.^[9]^ PLP-mediated enzymatic catalysis follows the recruitment of PLP from the environment, especially through interactions outside the active site.^[10]^ However, the precise mechanism by which enzymes recruit PLP remains unclear.

Amino acids, as prototypical substrates of PLP-dependent enzymes, likely play crucial roles in prebiotic chemistry.^[11,12]^ In the context of early Earth, amino acids might have been synthesized, transformed and degraded through nonenzymatic reactions under plausible primordial conditions.^[13–15]^ These transformations might have been mediated by PLP-like molecules, which are thermodynamically favorable and readily formed.^[11,16]^ During molecular evolution, PLP is thought to have initially associated with RNA (RNA-PLP) and later evolved to function within protein scaffolds, enabling greater diversity and efficiency of chemical reactions.^[11]^ The recruitment of PLP by enzymes would have conferred a significant selective advantage by accelerating amino acid turnover, an essential process for cellular growth and adaptation. Notably, enzymatic PLP catalysis proceeds 10^7^ to 10^9^ times faster than PLP catalysis alone, underscoring the importance of PLP capture as a rate-limiting and evolutionarily critical step in metabolism.^[15,17,18]^ Understanding how enzymes capture PLP is therefore essential for elucidating cofactor-driven molecular evolution and guiding enzyme engineering strategies.

Methionine γ-lyase (MGL), a representative PLP-dependent enzyme, catalyzes the γ-elimination of L-methionine (L-Met), yielding α-ketobutyric acid, methanethiol and ammonia. MGLs are members of the cystathionine γ-lyase family and contribute to the production of volatile organosulfur compounds, with implications for cardiovascular health, evolution of life and flavor biosynthesis.^[19,20]^ MGLs exhibit therapeutic significance by depleting L-Met in biological systems, thereby inhibiting cancer progression and promoting healthspan and longevity.^[21–23]^ Although PLP is required throughout the methionine cleavage reaction, the mechanism by which MGLs recruit PLP prior to catalysis remains unknown.^[8]^

In this study, we investigated the mechanism of PLP recruitment by MGL from *Yarrowia lipolytica* (yMGL, KEGG ID: YALI0C22088g), a strain noted for its robust L-Met degradation capacity.^[20]^ We solved the crystal structure of yMGL in complex with the substrate L-Met and cofactor PLP, elucidated the molecular interactions between PLP and yMGL *in vitro*, and demonstrated the *in vivo* relevance of this interaction for L-Met catabolism. Through a combination of biochemical and genetic characterization, supported by structural modeling, we identified the C-terminal region as a key determinant of PLP binding specificity. Furthermore, site-directed mutagenesis, PLP binding affinity assays together with conservation analysis revealed the conserved residue critical for PLP recognition across MGLs. These findings provide mechanistic insight into PLP recruitment and highlight evolutionary adaptations of PLP-dependent enzymes that enhance PLP specificity and catalytic efficiency. Our results deepen understanding of PLP-enzyme interactions and provide a framework for the rational engineering of cofactor-dependent enzymes.

## Results and Discussion

### The structure of the yMGL-L-Met-PLP complex

yMGL belongs to a newly identified subgroup of cystathionine γ-lyases and is phylogenetically distinct from the other structurally characterized or functionally annotated family members (Figure 1a).^[20]^ The X-ray crystal structure of yMGL in complex with the cofactor PLP and substrate L-Met was determined (PDB ID: 9UUW) (Figure 1b-f and Table S1). The enzyme adopts a dimeric architecture, with the active site formed at the interface between adjacent monomers (Figure 1b). Each monomer consists of three domains: an N-terminal domain (residues 7-63), a PLP-binding domain (residues 64-261) and a C-terminal domain (residues 262-379) (Figure 1b). At the active site of chain A, PLP is covalently linked to the ε-amino group of K204, forming a Schiff base; the phosphate group of PLP forms hydrogen bonds with G91 and T203; the carboxylate group of L-Met forms salt bridges with R344 and R357, and its amino group forms a hydrogen bond with T332 (Figure 1c,d). At the active site of chain B, PLP is likewise linked to K204 via a Schiff base (Figure 1e,f). The PLP phosphate interacts via hydrogen bonds with Y61 (A chain), L92 and T203 and forms a salt bridge with R63 (A chain); a π–π stacking interaction is observed between the PLP aromatic ring and Y113; L-Met is stabilized through salt bridges with R334 and R357 (Figure 1e).

**Figure 1.**
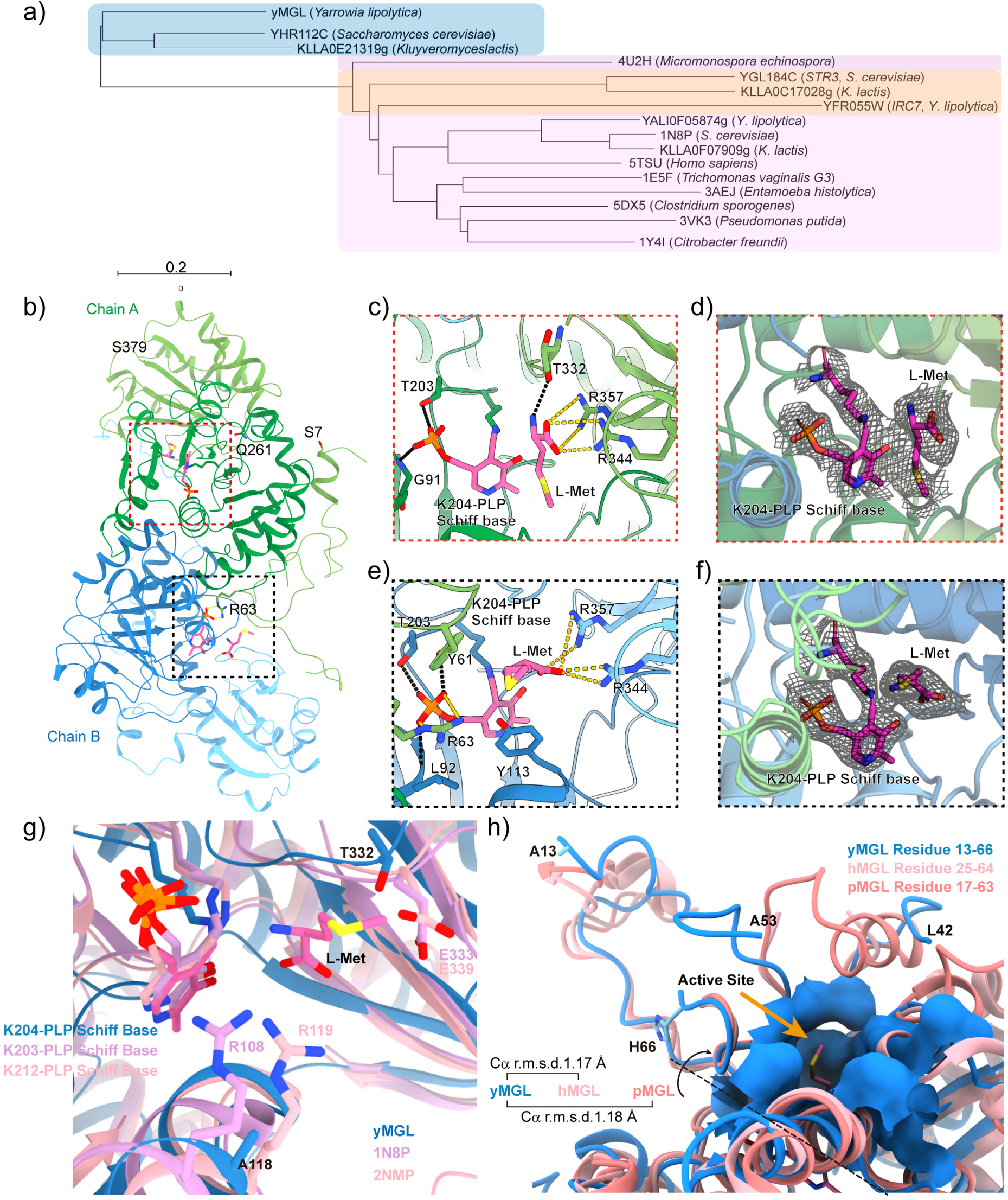
Phylogenetic and structural features of yMGL. (a) Phylogenetic relationships of MGLs from eukaryotes (*Homo sapiens*, *Trichomonas vaginalis*, *Entamoeba histolytica*, *Saccharomyces cerevisiae*, *Kluyveromyces lactis*, and *Yarrowia lipolytica*) and prokaryotes (*Pseudomonas putida*, *Citrobacter freundii*, *Clostridium sporogenes*, and *Micromonospora echinospora*). yMGL, representing the newly identified cystathionine γ-lyase group, is shaded in blue, and the other MGLs belonging to the cystathionine γ-lyase family are shaded in light purple. Cystathionine β-lyases from yeast are shaded in ochre. (b) Crystal structure of yMGL. The active sites exist at the interface of the two monomers. N-terminal domain, residues 7-63; PLP-binding domain, residues 64-261 (darker colors); C-terminal domain, residues 262-379. (c) A Schiff base was formed by PLP and K204 at the active site of chain A. L-Met was surrounded by the residues T332, R344 and R357. (d) Section of the simulated-annealing 2F_O_-F_C_ composite omit map (black mesh) contoured at the 0.7σ level at the active site of chain A. (e) A Schiff base was formed by the catalytic residues K204 and PLP at the active site of chain B. L-Met was surrounded by the residues Y61, Y113, R344 and R357. (f) Section of the simulated-annealing 2F_O_-F_C_ composite omit map (black mesh) contoured at the 1.0 σ level at the active site of chain B. The electron density for PLP in chain B is better resolved than that in chain A, likely because the absence of loop 53-64 is inferred to be involved in the interaction with PLP in chain B and its presence in chain A. (g) Comparison of the active sites in yMGL, *Saccharomyces cerevisiae* cystathione γ-lyase (PDB ID: 1N8P) and *Homo sapiens* cystathione γ-lyase (PDB ID: 2NMP, 2NMP was engineered to produce 5TSU, which contains the substitutions E59N, R119L, and E339V). Structural alignment of yMGL with 1N8P and 2NMP indicates that A118 and T332 (dodger blue) in yMGL, the charged residues R108 and E333 (plum) in 1N8P, and R119 and E339 (light pink) in 2NMP may dictate substrate selectivity between L-Met and L-cystathionine. (h) yMGL contains extended loops. Structural alignment of yMGL with hMGL and pMGL indicates that the active-site lid of yMGL comprises an extended loop segment, whereas the corresponding region in hMGL and pMGL is formed by multiple α-helices.

In yMGL and cystathionine γ-lyases (PDB IDs: 1N8P and 2NMP), PLP forms the conserved Schiff-base conformation with catalytic residue Lys (Figure 1g). In yMGL, the uncharged residues A118/T332 displace the conserved Arg/Glu residues that control the cystathionine selectivity in cystathionine γ-lyases, thereby conferring specific selectivity for L-Met (Figure 1g). The dimeric structure of yMGL resembles those of hMGL and pMGL, with Cα r.m.s.d. values of 1.17 Å and 1.18 Å, respectively (Figure 1h). Notably, yMGL contains an extended loop adjacent to the active-site lid, whereas the corresponding regions in the thermophilic enzymes hMGL and pMGL comprise several α-helices (Figure 1h). yMGL exhibits cold tolerance (Figure S1), and this extended loop may represent a structural adaptation that facilitates activity at low temperatures.^[24]^

The predicted entrance and active site of yMGL consists of a total of 33 amino acid residues (Figure 2a). Through alanine (Ala) scanning mutagenesis combined with L-Met degradation assays, we identified 22 key amino acid residues, which are highlighted in the spatial and primary structure of yMGL (Figure 2b-e). Notably, the majority of these essential residues, spanning from A28 to S244, are located within the N-terminal domain and PLP-binding domain (Figures 1b and 2a,e), with only one, S333, residing in the C-terminal domain (Figure 2a,e). These observations led us to further investigate the functional role of the C-terminal domain.

**Figure 2.**
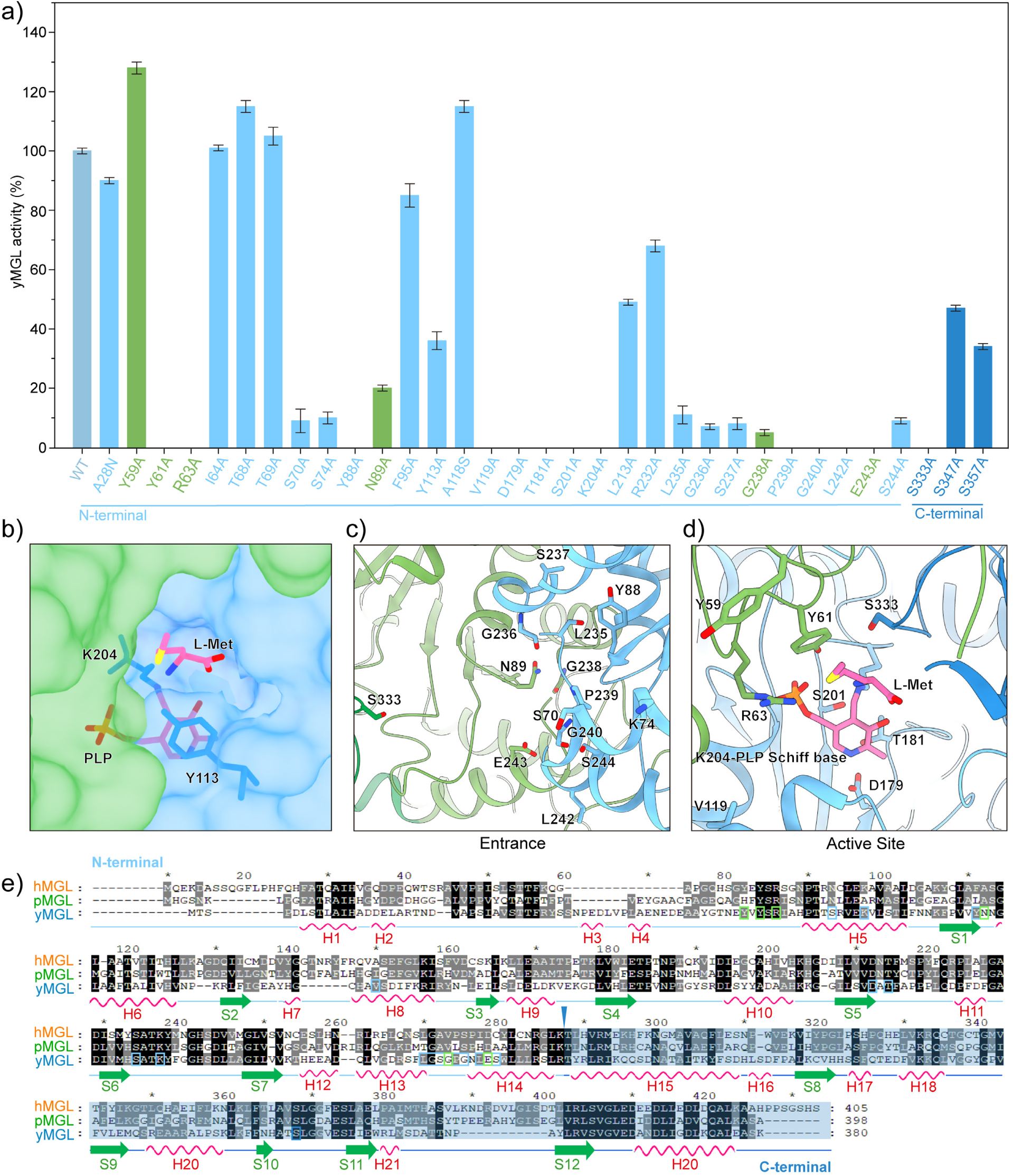
Key residues in yMGL. (a) Identification of key amino acid residues in yMGL. The amino acid residues in the entrance and active sites were mutated to Ala, and their activity was subsequently determined to identify the key amino acid residues. Only one key amino acid residue (S333) is located in the C-terminal region, which is shown with a deeper color. (b) Overview of the entrance and active sites of yMGL. (c,d) The key amino acid residues constituting the entrance (c) and active sites (d). These key residues are marked in color corresponding to the monomers of yMGL and shown in stick style. (e) Aligning the amino acid sequence of yMGL with pMGL (PDB ID: 2O7C) and hMGL (PDB ID: 5TSU). Identical amino acid residues are in black, and similar amino acid residues are in gray. Dashes represent gaps for improving the alignment. Secondary structure information was obtained from yMGL and presented as a ribbon diagram (H: Helix; S: Sheet). The truncated C-terminal domain is shaded, and the cleavage site is marked with a triangular wedge.

### Truncation of the C-terminal domain of yMGL

To investigate the functional role of the C-terminal domain in yMGL, we generated a truncated variant, yMGL-ΔCTD, by deleting residues 253-380 (Figures 2e and 3a). The theoretical molecular weights of yMGL and yMGL-ΔCTD as monomers are 41.93 kDa and 27.77 kDa, respectively, which is consistent with the results of the SDS-PAGE analysis (Figure 3b,c). Size exclusion chromatography revealed that both proteins eluted as single peaks, corresponding to apparent native molecular weights of ∼150 kDa for yMGL and 157 kDa for yMGL-ΔCTD (Figure 3d and Figure S2a). Dynamic light scattering analysis further revealed that yMGL-ΔCTD exhibited a larger hydrodynamic radius than yMGL did (Figure 3e). These data suggested that C-terminal truncation altered the quaternary structure from a homotetramer to a homohexamer at pH 8.0. This structural shift was accompanied by changes in the circular dichroism (CD) spectra (Figure S2b). More importantly, compared with full-length yMGL, yMGL-ΔCTD exhibited a 7.20-fold increase in L-Met degradation activity at optimal enzymatic reaction temperature (Figure 3f). However, the activity of yMGL-ΔCTD decreased more sharply at lower temperatures (Figures S1a and S2c), indicating reduced thermodynamic and kinetic stability (Figures S2d-f). The kinetic parameters of yMGL and yMGL-ΔCTD were determined using L-Met as the substrate under saturating PLP concentrations (Table 1 and Figure S3a,b). The catalytic efficiency of the yMGL-ΔCTD was 33.33% higher than that of the wild-type enzyme, which is closely associated with an enhanced binding ability for the L-Met (Table 1 and Figure S3a,b). However, when the L-Met concentration was maintained at saturation and the PLP concentration was varied, the catalytic efficiency of yMGL-ΔCTD decreased by 10.65-fold relative to that of yMGL (Table 1 and Figure S3c,d). These findings suggest that the C-terminal domain contributes to the specificity of yMGL for PLP.

**Table 1.**
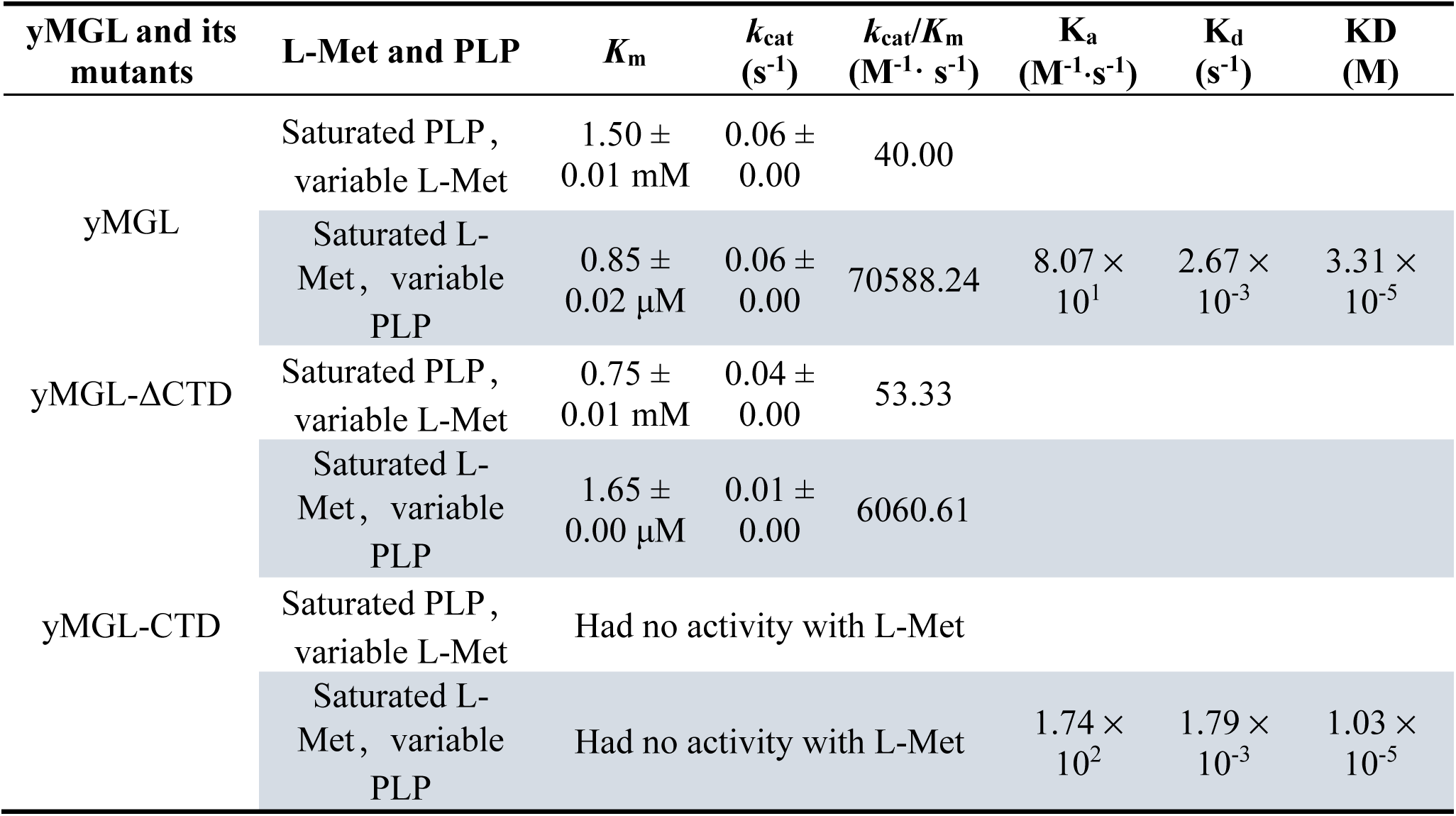
The parameters for kinetics and affinity.

| yMGL and its mutants | L-Met and PLP | $K_m$ | $k_{cat}$<br>(s<sup>-1</sup>) | $k_{cat}/K_m$<br>(M<sup>-1</sup>·s<sup>-1</sup>) | $K_a$<br>(M<sup>-1</sup>·s<sup>-1</sup>) | $K_d$<br>(s<sup>-1</sup>) | KD<br>(M) |
| --- | --- | --- | --- | --- | --- | --- | --- |
| yMGL | Saturated PLP,<br>variable L-Met | 1.50 ±<br>0.01 mM | 0.06 ±<br>0.00 | 40.00 |  |  |  |
|  | Saturated L-Met,<br>variable PLP | 0.85 ±<br>0.02 μM | 0.06 ±<br>0.00 | 70588.24 | 8.07 ×<br>10<sup>1</sup> | 2.67 ×<br>10<sup>-3</sup> | 3.31 ×<br>10<sup>-5</sup> |
| yMGL-ΔCTD | Saturated PLP,<br>variable L-Met | 0.75 ±<br>0.01 mM | 0.04 ±<br>0.00 | 53.33 |  |  |  |
|  | Saturated L-Met,<br>variable PLP | 1.65 ±<br>0.00 μM | 0.01 ±<br>0.00 | 6060.61 |  |  |  |
| yMGL-CTD | Saturated PLP,<br>variable L-Met | Had no activity with L-Met |  |  |  |  |  |
|  | Saturated L-Met,<br>variable PLP | Had no activity with L-Met |  |  | 1.74 ×<br>10<sup>2</sup> | 1.79 ×<br>10<sup>-3</sup> | 1.03 ×<br>10<sup>-5</sup> |

**Figure 3.**
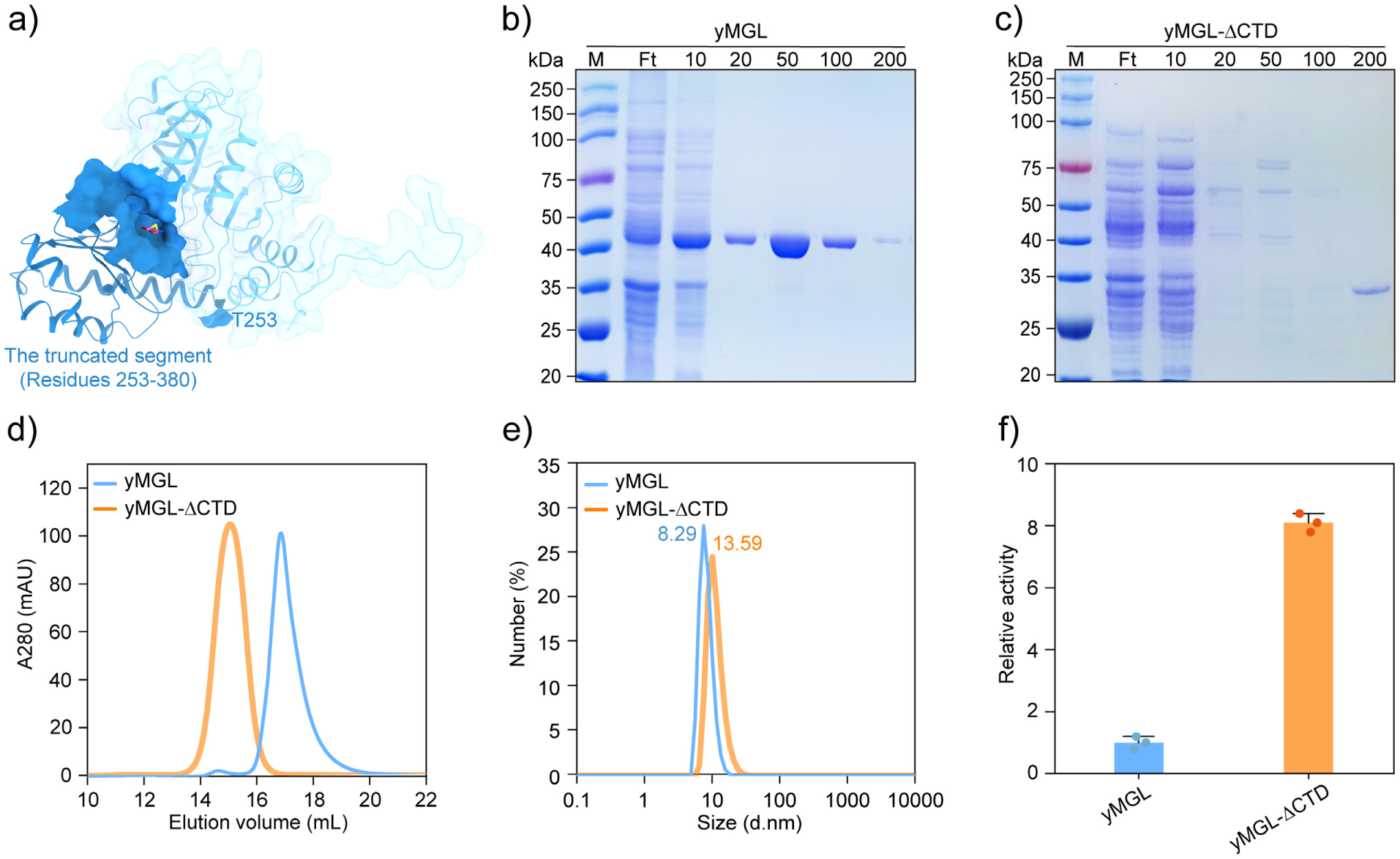
C-terminal truncation increases the molecular weight and L-Met lysis activity of multimers. (a) Truncation of the C-terminal region of yMGL. (b,c) SDS-PAGE analysis of yMGL (b) and its truncation mutant yMGL-ΔCTD (c). The theoretical monomer molecular weights are 41.93 kDa for yMGL and 27.77 kDa for yMGL-ΔCTD, consistent with the SDS-PAGE results. (d) Purification of yMGL and yMGL-ΔCTD by size exclusion chromatography. The native molecular weights are ∼150 kDa for yMGL and ∼157 kDa for yMGL-ΔCTD (Figure S2a). In solution, yMGL exists as a homotetramer, whereas yMGL-ΔCTD exists as a homohexamer. (e) Dynamic light scattering analysis of yMGL and yMGL-ΔCTD. yMGL-ΔCTD has a larger hydrodynamic size than yMGL. (f) yMGL-ΔCTD exhibited higher L-Met lyase activity than yMGL. The activity of yMGL at 20 °C was 299.06 ± 1.77 nmol·min^-1^· mg recombinant protein^-1^ and was normalized to 1.0.

### Functional analysis of the C-terminal domain of yMGL *in vitro* and *in vivo*

Binding affinity, a key indicator of substrate specificity in enzymes, was used to assess changes in specificity toward PLP between yMGL and the isolated C-terminal domain (yMGL-CTD).^[25–27]^ As shown in Table 1, the equilibrium dissociation constant (KD), reflecting the PLP-binding affinity, was 3.31 × 10^-5^ M for yMGL (Figure 4a). The KD value for yMGL-CTD was 1.03 × 10^-5^ M (Figure 4b).

**Figure 4.**
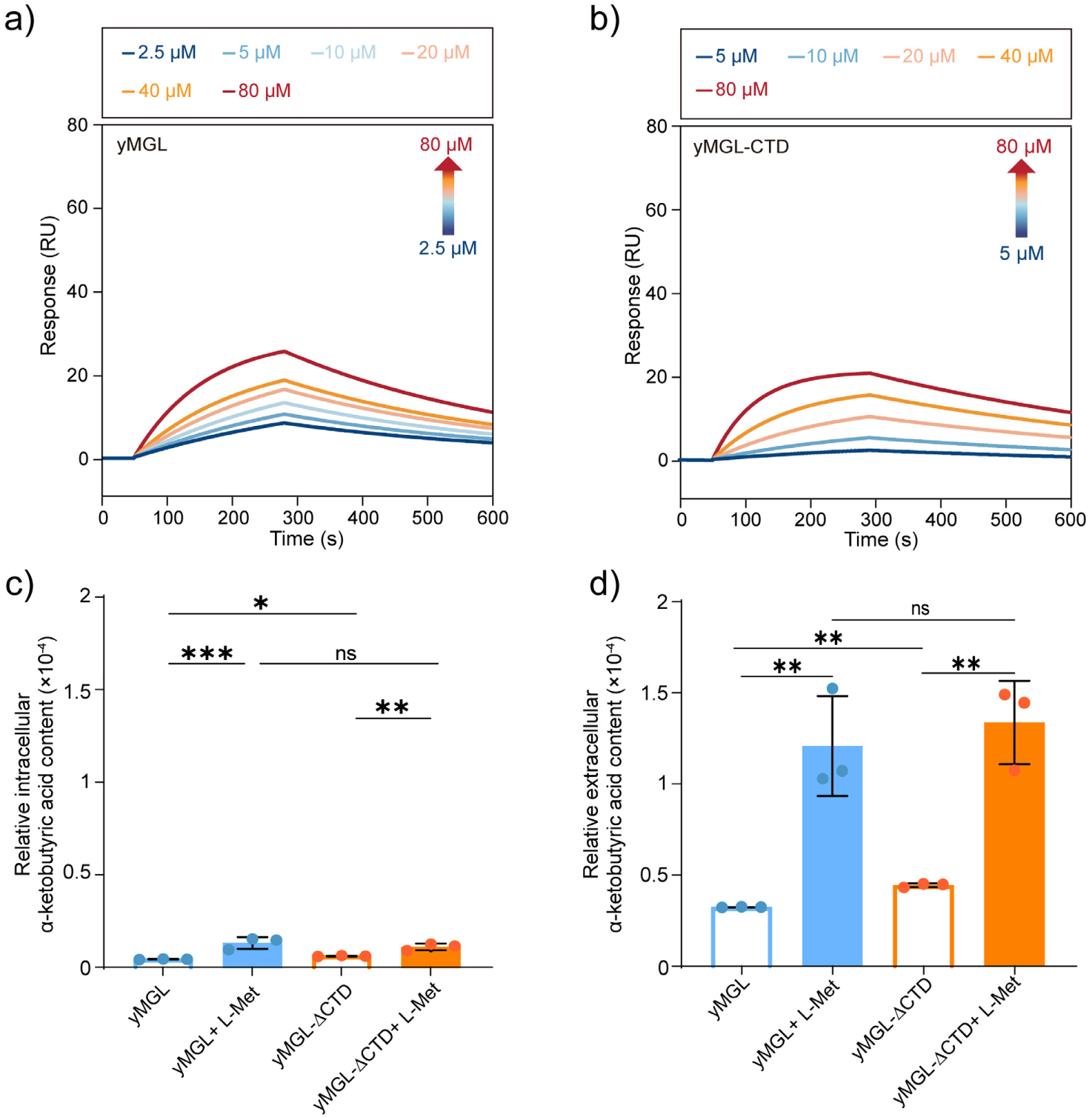
The C-terminal region specifically binds PLP and its truncation impacts L-Met degradation in *Y. lipolytica*. (a) yMGL shows strong binding affinity for PLP (K_a_=8.07 × 10^1^ M^-1^·s^-1^, K_d_=2.67 × 10^-3^ s^-1^, KD=3.31 × 10^-5^ M). (b) The binding affinity between PLP and isolated yMGL-CTD. PLP-binding affinity of yMGL-CTD (K_a_=1.74 × 10^2^ M^-1^·s^-1^, K_d_=1.79 × 10^-3^ s^-1^, KD=1.03 × 10^-5^ M) is comparable to that of full-length yMGL. (c,d) The effect of C-terminal domain deletion on the intracellular (c) and extracellular (d) accumulation of α-ketobutyric acid. yMGL-ΔCTD accumulated higher levels of intracellular and extracellular α-ketobutyric acid than yMGL. This observation may be attributable to sufficient intracellular PLP, which enhances the lyase activity of yMGL-ΔCTD (Figure 3f). Supplementation with L-Met markedly increased extracellular α-ketobutyric acid accumulation, consistent with active excretion.[30] Statistical analyses were performed using GraphPad Prism 8.3.0 Software, *: P <0.05; **: P<0.01; ***: P<0.001; ns: not significant.

These values are of the same order of magnitude and indicate that the C-terminal domain substantially contributes to yMGL’s specific affinity for PLP.

To assess the effect of C-terminal domain implicated in PLP specificity on L-Met degradation *in vivo*, we deleted the sequence encoding the C-terminal domain by using a CRISPR-Cas9 system integrated with homologous recombination technology.^[28,29]^ The deletion genotypes were confirmed by observing the reduction in the PCR product size and sequencing the PCR products (Figure S4 and Table S2). The deletion of the C-terminal domain in yMGL increased intracellular and extracellular α-ketobutyric acid levels (Figure 4c,d). C-terminal domain truncation altered the protein’s oligomeric assembly (Figures 3d,e), and adequate intracellular PLP (both strains were cultured in minimal medium supplemented with 4.86 μM PLP to support growth) levels may confer higher L-Met lyase activity on yMGL-ΔCTD (Figure 3f). The supplementation of L-Met markedly increased α-ketobutyric acid accumulation in both strains and resulted in extracellular α-ketobutyric acid levels exceeding intracellular levels (Figure 4c,d), likely reflecting active excretion to alleviate the compound’s cytotoxicity.^[30]^

### Key amino acid residues in the C-terminal region that determine the specific action of PLP

The homotetrameric structure of yMGL within the enzymatic system was modeled by assembling two dimeric units (Figure 5a). The C-terminal domains extend across the active site and lie above its entrance (Figure 5a-c). Residues R255, K270 and S333 in the C-terminal domain were predicted to interact with PLP (Figure 5a). However, only the S333A mutant (yMGL-S333A) reduced PLP-binding affinity, by 27.54-fold (Figures 4a and 5d). Furthermore, the S333A mutant exhibited a 3.4 °C decrease in melting temperature (an indicator reflecting the substrate specificity) (Figure 5e,f). Together, these data suggested that S333 determines the specific interaction between the C-terminal domain and PLP. Either the S333 residue of the active-site monomer or the S333 residue (the adjacent monomer) lying near the active-site entrance is inferred to interact with PLP (Figure 5a). Notably, this Ser residue is highly conserved among MGL homologs (Figure 2e). In hMGL, the S340A mutation, which is positioned at the active-site entrance, markedly reduced PLP-binding affinity and specificity, and the S340A mutant showed a 7.6 °C decrease in melting temperature (Figure 6). We therefore conclued that the Ser at the entrance is critical for PLP recruitment by yMGL.

**Figure 5.**
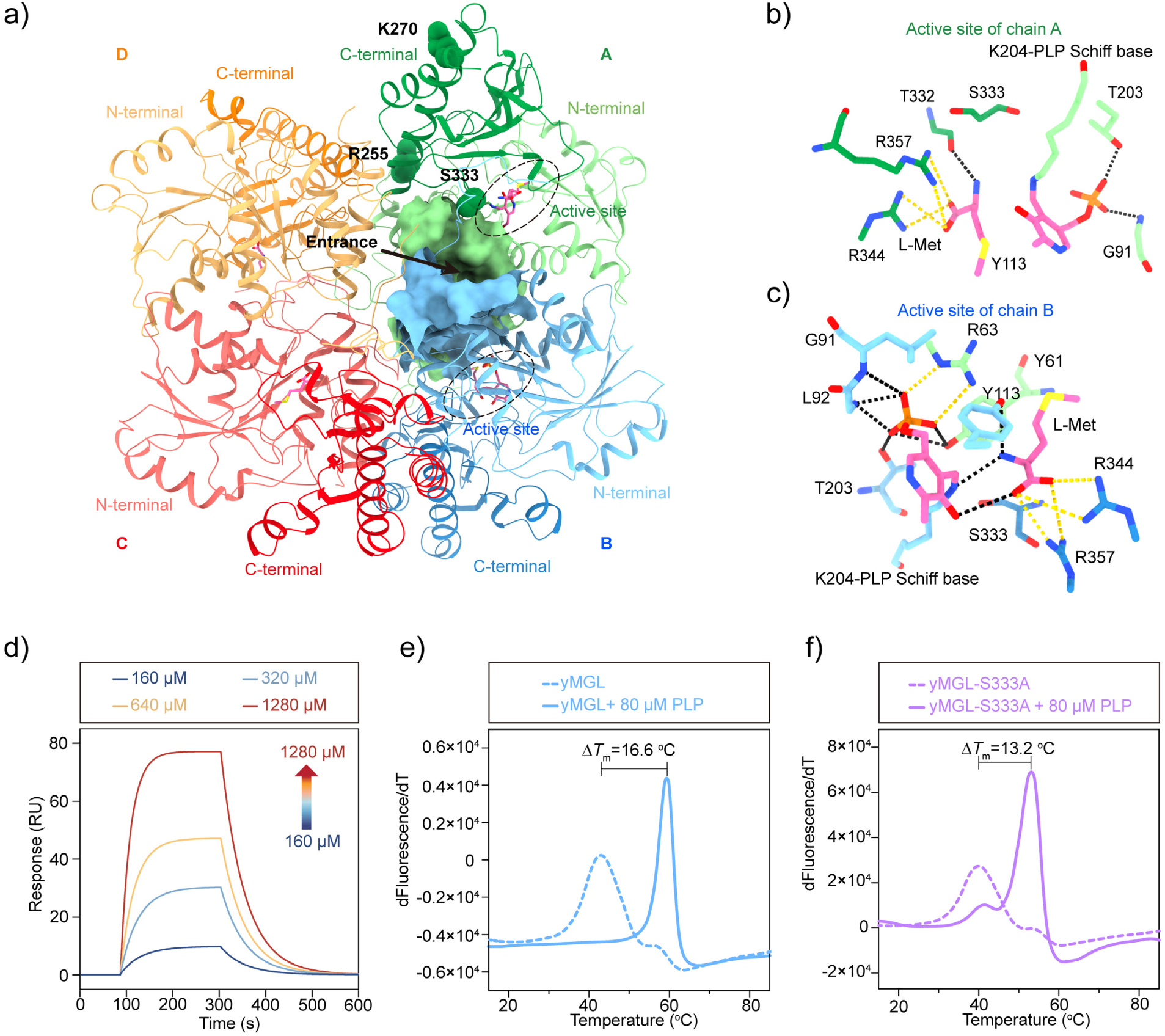
Identification of key C-terminal domain residues that specifically interact with PLP. (a) Screening the amino acid residues with the potential to specifically interact with PLP. R255, K270, and S333 were predicted to interact with PLP and are displayed as ball-and-stick models. (b, c) S333 is located at the active site (b) or near the entrance (c). At the active site of chain A (b), PLP forms a Schiff base with K204 and has no direct interaction with S333; at the active site of chain B (c), S333 similarly lacks direct contact with PLP. (d) The PLP-binding affinity for yMGL-S333A. Mutating S333 into Ala in yMGL sharply decreases its specificity for PLP by 27.54-fold (K_a_=2.08 × 10^1^ M^-1^·s^-1^, K_d_=1.96 × 10^-2^ s^-1^, KD=9.42 × 10^-4^ M). (e,f) The binding affinity toward PLP was was further evaluated via a thermal shift assay. PLP addition increased the melting temperature (*T*_m_) of yMGL (e), whereas the S333A mutation (f) markedly decreased *T*_m_ relative to wild type, consistent with reduced PLP specificity.

**Figure 6.**
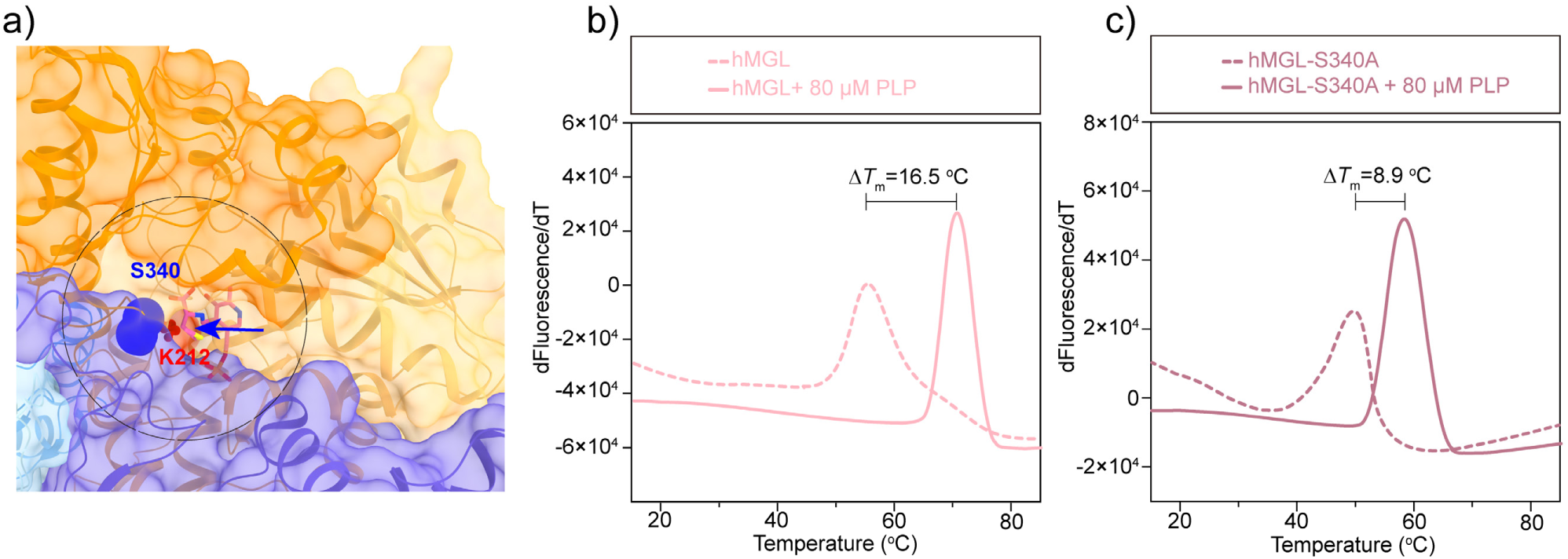
Functional identification of S340 in hMGL. (a) S340 situated at the proximity of the entrance of the hMGL. S340 in hMGL is conserved with S333 in yMGL and is shown as a blue ball model. The entrance and active sites in hMGL are predicted to occur at the interface of the dimer; K212, the inferred catalytic residue, is shown as a red ball-and-stick model. The C-terminal domains of each monomer are depicted in darker shades. (b,c) Binding affinity toward PLP, assessed by thermal shift assay. Addition of PLP increased the melting temperature (*T*_m_) of hMGL (b), whereas substitution of S340 with Ala (c) markedly reduced *T*_m_ relative to wild type, indicating decreased PLP specificity.

AlphaFold3 modeling of the yMGL-ΔCTD variant indicated that the active sites become surface-exposed and that, at the active sites, PLP forms hydrogen bonds with T56, E58, V60, and S233 (Figure S5). Thus, yMGL-ΔCTD is expected to recruit PLP from the environment more easily. While its PLP-binding ability was obviously lower than that of yMGL (Table 1 and Figure S3c,d). yMGL and yMGL-CTD displayed comparable affinities for PLP (Table 1 and Figure 4a,b). These results prompted re-evaluation of structural features of the C-terminal domain that may specifically enhance the Ser-PLP interaction. The conserved Ser, locating within a extended loop, positioned near the active-site entrance of yMGL and hMGL (Figures. 5a and 6a), where they may act as a PLP-trapping module that facilitates the Ser–PLP interaction and guides PLP into the hydrophobic pocket, thereby enhancing catalytic efficiency.

To investigate how the MGLs evolved to recruit PLP, we utilized an ancestral sequence reconstruction strategy to obtain putative ancestral homologs. Ten different ancestral nodes were inferred from the phylogenetic tree of the yMGL subfamily (Figure 7a). Sequence analysis and structural alignment revealed loop regions surrounding the conserved Ser in the C-terminal domain of yMGL, its reconstructed ancestors and other MGLs (Figure 7b-d), consistent with a PLP-trapping mechanism for cofactor recruitment. yMGL exhibited cystathionine γ-synthase activity and shared ancestral homologs with cystathionine γ-synthases (Figures 7a and S6), suggesting that it retains the broad substrate promiscuity of its ancestors. Truncating the C-terminal domain of yMGL altered the overall structural stability and reduced PLP specificity (Figures 4a,b, S2d-f, S3c,d and Table 1), implying that this domain may have evolved to mediate high-affinity PLP recruitment, a key step for efficient enzymatic L-Met degradation. The inherent adaptability of these loops in yMGL, its ancestral homologs and other MGLs makes them promising targets for protein engineering. Indeed, loop engineering has been shown to expand the functional landscape of enzymes by modulating their activity, specificity, and selectivity, and may enable the rational design of enzymes with enhanced cofactor-binding capabilities at distal sites within the protein.^[31–33]^

**Figure 7.**
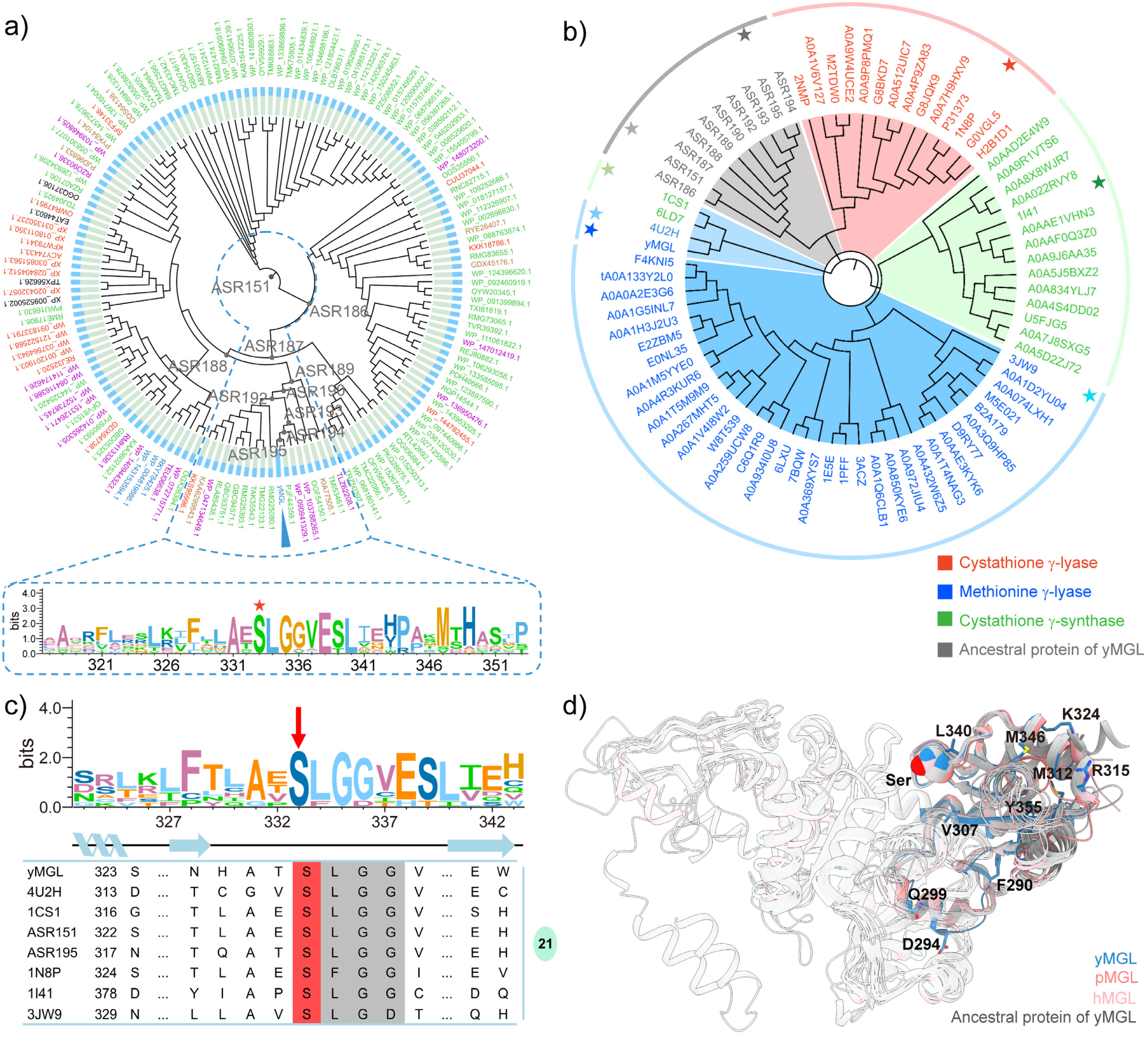
Ancestral reconstruction of yMGL reveals conservation of Ser residue. (a) The Ser residue is conserved in yMGL and its ancestral homologs. Sequences of cystathionine γ-synthases, cystathionine γ-lyases, aminotransferases, cystathionine β-lyases, hypothetical proteins and MGLs (shown in green, red, purple, brown, black and blue, respectively) dependent on PLP with accession numbers were obtained from the FireProt server for phylogenetic analysis with yMGL. A phylogenetic tree was then constructed to infer ancestral homologs, which included ASR195, ASR194, ASR193, ASR192, ASR190, ASR189, ASR188, ASR187, ASR186, and ASR151. (b) Phylogenetic analysis of yMGL, its ancestral homologs, other MGLs, cystathionine γ-synthases and cystathionine γ-lyases. yMGL clusters with the MGL (PDB ID: 4U2H) and the cystathionine γ-synthases (PDB IDs: 6LD7 and 1CS1), forming a distinct phylogenetic tree branch. This suggests that yMGL may have retained the substrate promiscuity characteristic of its ancestors. (c) Multiple sequence alignment of eight representative proteins. The Ser residue (indicated by a red arrow) is strictly conserved among the eight proteins marked with asterisks in Figure 7b, including yMGL, 4U2H, 1CS1, ASR151, ASR195, 1N8P, 1I41 and 3JW9. (d) Structural superposition of yMGL with its ten ancestral homologs (shown in gray), hMGL, and pMGL. The conserved Ser residue (shown in ball representation) is located within a loop-rich region of the C-terminal domain. The C-terminal domains are color-coded, and the loops encompass residues 290-294, 299-307, 312-315, 324-340, and 346-355 (yMGL numbering).

## Conclusion

yMGL represents a newly identified subgroup within the cystathionine γ-lyase family. In this study, we report its crystal structure (PDB ID: 9UUW) and show that the C-terminal domain-located outside the canonical PLP-binding domain-determines specific PLP affinity and is required for L-Met degradation. A Ser residue in this C-terminal domain is conserved in yMGL, its reconstructed ancestral homologs, other MGLs, and mediates the Ser-PLP interaction in both yMGL and hMGL. We infer that MGLs have evolved a PLP-trapping mechanism in which a conserved Ser promotes selective PLP recruitment at a structural motif near the active-site entrance. These findings advance our understanding of PLP recognition and recruitment by enzymes and provide a mechanistic framework for the molecular evolution of PLP-dependent enzymes. Moreover, they offer a rational basis for engineering enzymes with altered or enhanced cofactor specificity.

## Method

### Crystallization and structure determination of yMGL

yMGL was expressed in *E. coli* and purified by nickel affinity chromatography following the method described by Zhao *et al.* ^[20]^ and then yMGL was further purified by an anion charge column Resource Q (GE Lifesciences) and Superdex200 Increase 10/300GL gel-filtration column (GE Healthcare) following the method described by Jia.^[34]^ yMGL was incubated with 0.2 mM PLP and 1 mM L-Met for 30 min before being subjected to crystallization screening at 4 °C. The crystallization condition for the best diffracted crystal was 0.2 mol/L NaCl, 0.1 mol/L CAPS, pH 10.3, 20% PEG8000, and 11 mg/mL yMGL. Diffraction data were collected on beamline BL02U1 at the Shanghai Synchrotron Radiation Facility (SSRF). X-ray diffraction data were collected from a single crystal at 100 K using a monochromatic X-ray at a wavelength of 0.97918 Å. The data were processed and scaled by XDS (PMID: 20124692) and Aimless (PMID: 27275144). The structure of yMGL in complex with L-Met was solved by molecular replacement with the program Phaser (PMID: 19461840) using the model built by AlphaFold as the template. Several rounds of manual building by Coot (PMID: 15572765) and refinement by Phenix (PMID: 20124702) were carried out. The statistics of X-ray data collection and refinement are summarized in Appendix Table S1. Structure figures were prepared by ChimeraX.

### Mutagenesis of MGLs

The pET28a plasmid was used to express MGLs and their mutants. A Mut Express^®^ II Fast Mutagenesis Kit V2 (Vazyme, Nanjing, Jiangsu Province, China) was used to mutate amino acid residues by using the primers detailed in Table S2. The PCR amplification products were digested by *Dpn*I to remove the template DNA and then used directly in the recombination reaction. pET28a containing the genes encoding MGLs and their mutants was subsequently transformed into *E. coli* BL21 (DE3). Single colonies with the target mutation were inoculated into LB culture media supplemented with 50 μg/mL kanamycin and cultured overnight at 37 °C. After the optical density at 600 nm approached 0.8, 0.5 mM isopropyl thiogalactoside (IPTG) was added to the medium to induce the expression of MGLs and their mutants at 18 °C. After growing for 20 h, the cells were harvested by centrifugation, after which the MGLs and their mutants were purified.

### MGL activity determination

The purified MGLs were treated with phenylhydrazine hydrochloride to remove the naturally bound PLP following the protocol.^[35,36]^ MGL activity was assayed following the method described by Zhao *et al.*^[20]^ The concentration of L-Met (Shanghai Aladdin Biochemical Technology Co., Ltd., 99% pure) was 20 mM, and the concentration of PLP (TCI (Shanghai) Development Co., Ltd., over 98% pure) was 5 mM in a total volume of 1 mL of reaction mixture with 0.6 mg of the purified MGLs added. The inactivated enzymes (after boiling for 10 min) were used as the control. HPLC (Dionex UltiMate 3000, Thermo) with a Reprosil-Pur Basic C18 column (4.6 mm × 250 mm × 5 μm) from Dr Maisch GmbH (Germany) was used to analyze the decrease in the L-Met concentration and the production of α-ketobutyric acid. A 13%/87% acetonitrile/water mixture adjusted to pH 3.0 with H_3_PO_4_ was used as the mobile phase, and the detection wavelength was 230 nm. The kinetic stability (*t*_1/2_) and thermodynamic stability (*T*_m_) of yMGL and its mutant were measured following methods of Hu et al.^[23]^

### Determination of the kinetics, CD and binding affinity

The enzymatic degradation of L-Met was assessed in a final 1 mL volume containing various concentrations (0.1-50 mM) of L-Met, 5 μM PLP, 50 mM Tris-HCl buffer (pH 8.0) containing 0.6 mg/mL yMGL and its mutants. The reactions were performed at 20 °C for 10 min, after which 5 μL of H_3_PO_4_ (10 M) was added to the mixture to quench the reactions. To determine the kinetics of PLP, the PLP concentration was varied from 0.1 to 20 μM, and the L-Met concentration was fixed at 20 mM. The production of α-ketobutyric acid from L-Met was detected by reaction with MBTH following a method described previously.^[37]^

A Chirascan V100 Spectrometer (Applied Photophysics, Leatherhead, Surrey, UK) was used to measure the CD spectra of yMGL and its truncation mutant. Both were suspended in 50 mM Na_2_HPO_4_/NaH_2_PO_4_ (pH 8.0) buffer at equal molar concentrations, and then CD spectra were obtained from 190 to 280 nm (a step size of 1 nm, 0.5 s time-per-point, 1 nm bandwidth). After the baseline was corrected, each sample was scanned three times to obtain the spectrum.

The interaction between PLP and yMGL or its mutants was analyzed using a Nicoya OpenSPRTM system at 20 °C following the method described by Tan *et al.*^[38]^ yMGL or its mutants were immobilized on the sensor Chip NTA (Cat. No. SEN-AU-100-10-NTA). PLP was injected as an analyte at various concentrations, and PBS (pH 8.0) was used as the running buffer to determine the K_a_ (association rate), K_d_ (dissociation rate), and KD (equilibrium dissociation constant that reflects the affinity for substrates).

The PLP-binding affinities of yMGL, hMGL, and their variants were determined using a thermostability shift assay (TSA). For each sample, a 1 μg/μL protein solution was prepared in reaction buffer, and AGSYPRO Orange dye (5,000× stock; Accurate Biology Co.) was added to a final concentration of 5×. Measurements were performed in 96-well format on a QuantStudio 3 Real-Time PCR system (Thermo Fisher Scientific) using the x1 m3 detection channel, and fluorescence was recorded with QuantStudio Design & Analysis Software (v1.3.1). The temperature was increased at a rate of 0.05 °C/s from 15 °C to 85 °C, and the *T*_m_ was determined from the peak of the first derivative of the melt curve.

### Deletion of the C-terminal domain of yMGL in *Y. lipolytica*

The optimal gRNA sequence (20 bp) targeting yMGL was selected by using CHOPCHOP (https://chopchop.cbu.uib.no/), with W29 as the genetic background.^[28]^ The plasmid pCfB3405 was used in this study to construct vectors harboring a single gRNA expression cassette.^[29]^ The genomic DNA of W29 was used as a template for amplifying endogenous fragments that overlapped to generate seamlessly integrated fragments. To obtain a recombinant strain, the integrated fragments and specific gRNA plasmids were mixed and transformed into *Y. lipolytica* by using the lithium acetate-mediated yeast transformation method.^[29]^

*Y. lipolytica* was cultured in minimal medium (7.5 g/L (NH_4_)_2_SO_4_, 4.4 g/L KH_2_PO_4_, 0.5 g/L, MgSO_4_⋅7H_2_O, 20 g/L glucose, 2 mL/L micronutrient metals and 1 mL/L vitamins).^[39]^ *Y. lipolytica* cells and fermentation supernatant samples were analyzed via a Vanquish ultrahigh-performance liquid chromatography (UHPLC) system (Mercedes-Benz, Bremen, Germany) at 36 h. After the raw LC‒MS data were calibrated, the MS information was simultaneously matched with the metabolism public database HMDB and the Majorbio cloud platform (https://www.majorbio.com/) to obtain the metabolite information of α-ketobutyric acid.

### Molecular modeling and phylogenetic analysis of MGLs

A nonrooted molecular phylogenetic tree of the MGLs from *Homo sapiens*, *Trichomonas vaginalis*, *Entamoeba histolytica*, *Saccharomyces cerevisiae*, *Kluyveromyces lactis* and *Yarrowia lipolytica*, which are eukaryotes; *Pseudomonas putida*; *Citrobacter freundii*; *Clostridium sporogenes*; and *Micromonospora echinospora*, which are prokaryotes, was established through ClustalW multiple sequence alignment via the maximum likelihood method in MEGA X 10.0.4. The phylogenetic relationships of MGLs from yeast were analyzed via the fast minimum evolution method at the website (https://blast.ncbi.nlm.nih.gov/Blast.cgi). MGLs and their mutants in enzymatic reactions were modeled via AlphaFold3 structural prediction via a web server (https://alphafoldserver.com/) and visualized in ChimeraX.^[40,41]^ The entrance of yMGL was predicted via CAVER software (https://loschmidt.chemi.muni.cz/hotspotwizard/) and displayed by ChimeraX. AutoDock software was used to analyze the interaction between PLP and MGLs. The ancestral sequence of yMGL was reconstructed through the FireProt-ASR online service at the website (http://loschmidt.chemi.muni.cz/fireprotasr/). ^[42]^

## Acknowledgements

This work was supported by the National Key Research and Development Program of China (2022YFC2106100), the National Natural Science Foundation of China (Grant Nos. 31570054, 22377098 and 21807088), the Key Project of Hubei Provincial Department of Education (T2022011) and the State Key Laboratory of Microbial Technology Open Projects Fund (Project No. M2022-06). We thank the SSRF BL02U1 beam line for data collection and processing.

