## Supplement Information for "Methionine γ-lyase Traps Cofactor Pyridoxal-5′-phosphate through Specific Serine-Mediated Affinity"

### Table of content


**Figure S1.** Enzymatic characterization of yMGL ···································· 3

**Figure S2.** Enzymatic characterization of yMGL and yMGL-ΔCTD ···································· 4

**Figure S3.** Kinetic parameters for yMGL and yMGL-ΔCTD measured under saturating PLP or L-Met concentrations ···································· 5

**Figure S4.** The deletion of the sequence encoding the C-terminal domain of yMGL ···································· 6

**Figure S5.** Simulation of the interaction between PLP and yMGL-ΔCTD ···································· 7

**Figure S6.** yMGL functions as a cystathionine γ-synthase ···································· 8

**Table S1.** Data and refinement statistics for the structures of yMGL ···································· 9

**Table S2.** Primers used for constructing the yMGL mutants ···································· 10

**Figure S1.**

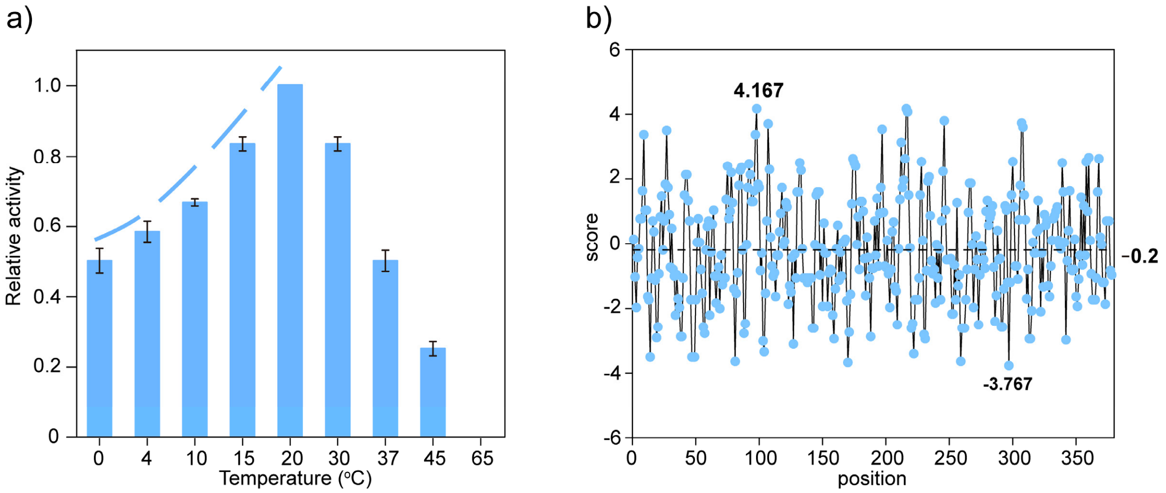


**Figure S1.** Enzymatic characterization of yMGL. (a) Activity of yMGL at different temperatures. The optimal reaction temperature was 20 °C, and 50% of the maximum activity was retained at 0 °C, indicating cold tolerance. (b) Hydrophobic analysis of yMGL. The hydrophobicity and hydrophilicity are represented by positive values and negative values, respectively. 0 represents neutrality. yMGL has a lower mean hydrophobicity value than pMGL and hMGL, both of which are thermophilic, consistent with its cold-adapted properties.[1,2]

**Figure S2.**

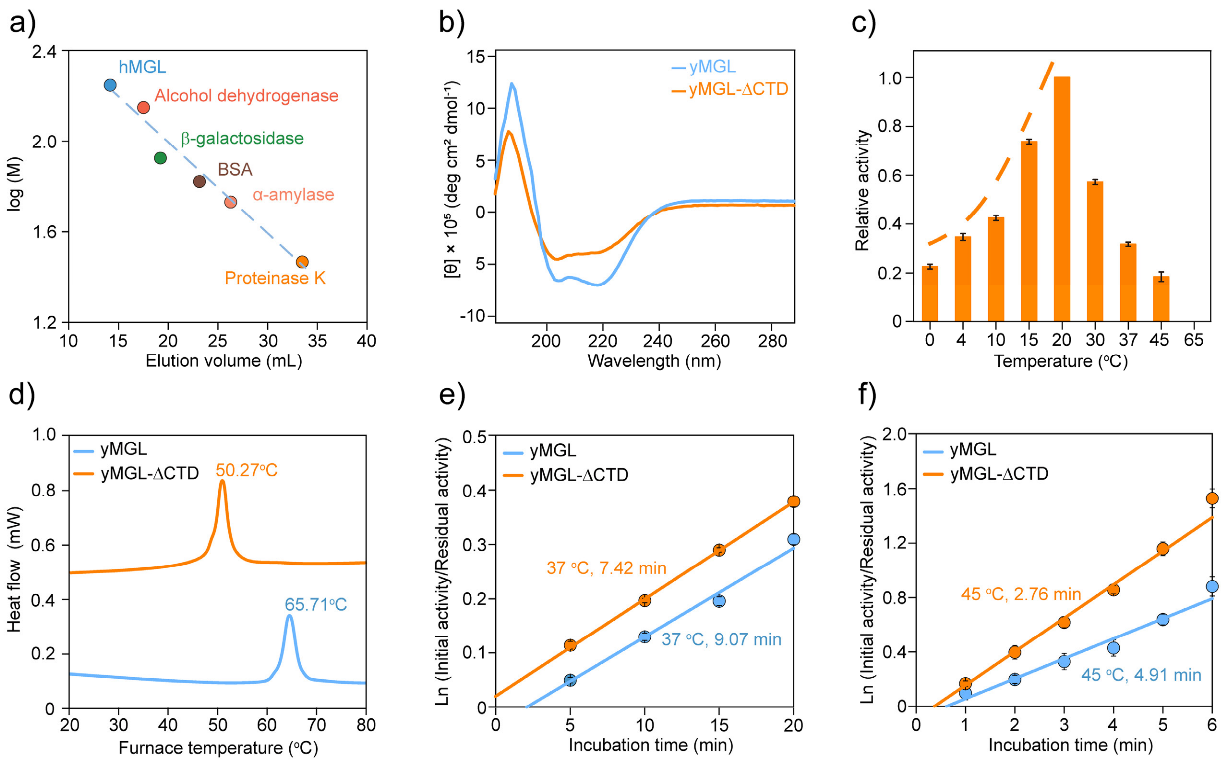


**Figure S2.** Enzymatic characterization of yMGL and yMGL-ΔCTD. (a) Calibration curves were used to determine the molecular weights of yMGL and its mutants. The molecular weights of proteinase K, α-amylase, BSA, β-galactosidase, alcohol dehydrogenase and hMGL are 29.3, 54.0, 66.4, 90.0, 141.0, and 176.8 kDa, respectively. (b) Truncation of the C-terminal domain changes the CD spectra of yMGL. (c) Activity of yMGL-ΔCTD at different temperatures. yMGL-ΔCTD exhibited maximum activity at 20 °C, with activity declining sharply at lower temperatures; at 0 °C, 22% of the maximum activity remained. (d) Thermodynamic stability of yMGL and yMGL-ΔCTD. $T_m$ values were determined by differential scanning calorimetry (DSC) and corresponded to 65.71 °C for yMGL and 50.27 °C for yMGL-ΔCTD. (e,f) Residual activities of yMGL and yMGL-ΔCTD. yMGL and yMGL-ΔCTD were incubated at 37 (e) and 45 °C (f), and their residual activities were measured at each point in incubation. The following kd values were calculated from the slope for yMGL: 0.017/min at 37 °C and 0.15/min at 45 °C; and for yMGL-ΔCTD: 0.018/min at 37 °C and 0.27/min at 45 °C.

**Figure S3.**

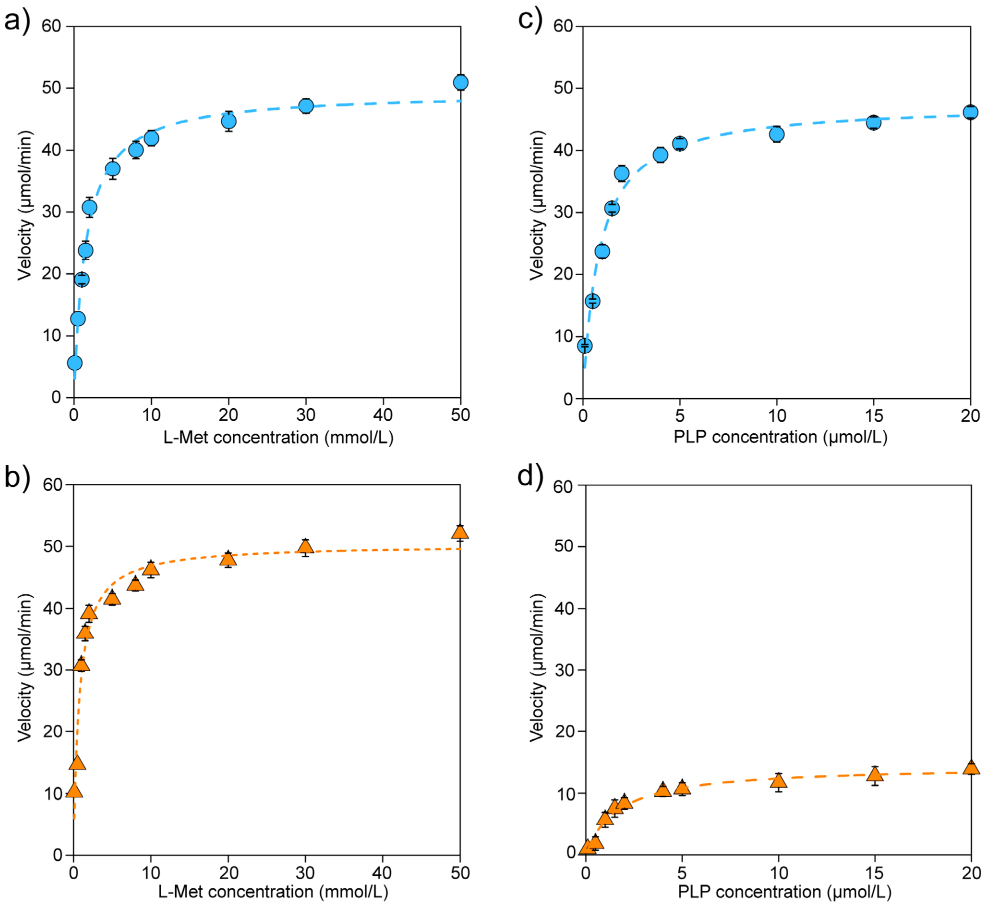


**Figure S3.** Kinetic parameters for yMGL and yMGL-ΔCTD measured under saturating PLP (a,b) or L-Met (c,d) concentrations.

**Figure S4.**

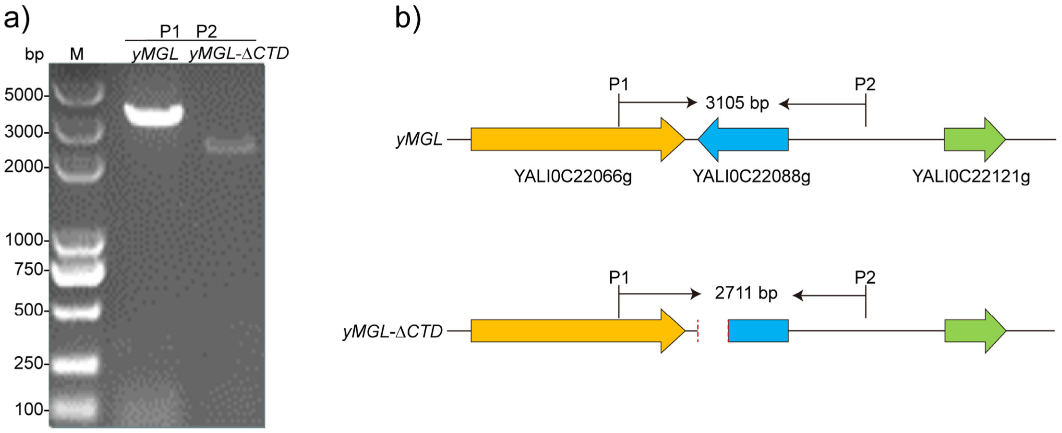


**Figure S4.** The deletion of the sequence encoding the C-terminal domain of yMGL. (a) Deletion of the sequence encoding the C-terminal domain of yMGL was confirmed by PCR. (b) The expected deletion of the sequence encoding the C-terminal domain of yMGL (yMGL-*ΔCTD*).

**Figure S5.**

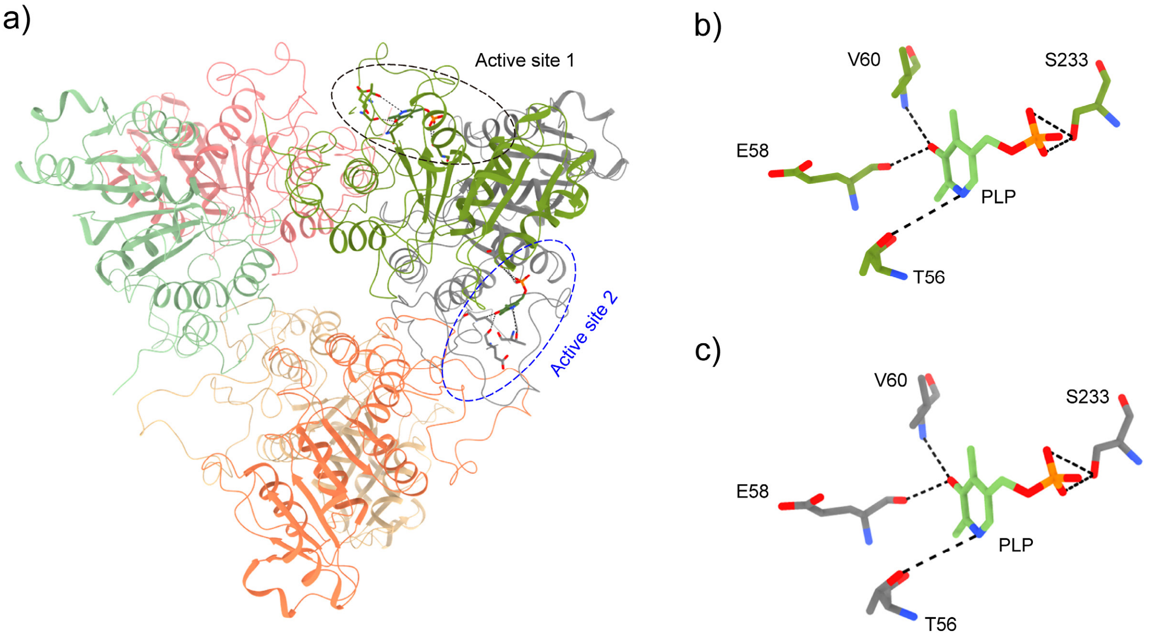


**Figure S5.** Simulation of the interaction between PLP and yMGL-ΔCTD. (a-c) Predicted model of the interaction between PLP and yMGL-ΔCTD (a). The predicted yMGL-ΔCTD is a hexamer, and PLP was docked at the two active sites that are present at the junction of two monomers and exposed on the surface. PLP interacts with residues at the two active sites via hydrogen bonding (b,c).

**Figure S6.**

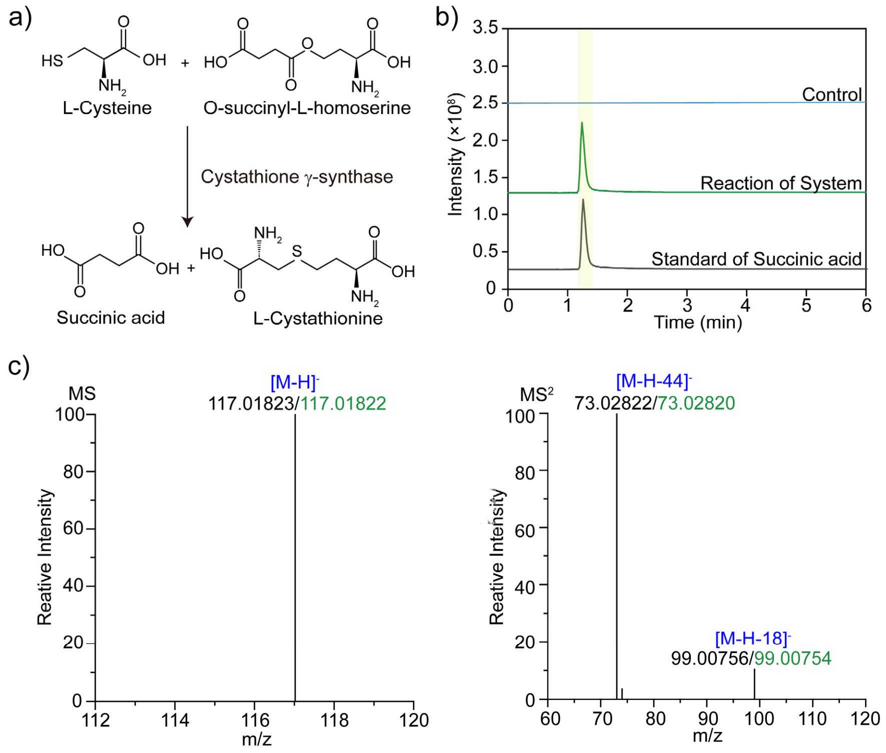


**Figure S6.** yMGL functions as a cystathionine γ-synthase. (a) Synthesis of L-cystathionine catalyzed by cystathionine γ-synthase.[3] (b-c) Succinic acid was produced in an enzymatic system containing L-cysteine and O-succinyl-L-homoserine as substrates (b) and confirmed by MS/MS (c). L-cystathionine was not detected, consistent with yMGL's ability to degrade L-cystathionine.[4]

**Table S1.** Data and refinement statistics for the structures of yMGL

| | yMGL |
|---|---|
| PDB ID | 9UUW |
| Data collection | |
| Wavelength | 0.97918 Å |
| Resolution range | 53.86 - 3.04 (3.149 - 3.04) |
| Space group | P 32 2 1 |
| Unit cell | |
| a, b, c (Å) | 166.054 166.054 81.314 |
| α, β, γ (°) | 90 90 120 |
| Total reflections | 520570 |
| Unique reflections | 25114 (2489) |
| Multiplicity | 20.7(21.4) |
| Completeness (%) | 99.35 (100.00) |
| Mean I/sigma(I) | 7.0 (1.9) |
| Wilson B-factor | 43.74 |
| R-merge | 0.942 (2.121) |
| R-meas | 0.966 (2.171) |
| R-pim | 0.211 (0.466) |
| CC1/2 | 0.966 (0.484) |
| Refinement | |
| Reflections used in refinement | 24958 (2489) |
| Reflections used for R-free | 1210 (130) |
| R-work | 0.2327 (0.3085) |
| R-free | 0.2662 (0.3374) |
| Number of non-hydrogen atoms | 5613 |
| macromolecules | 5613 |
| Protein residues | 716 |
| RMS (bonds) (Å) | 0.007 |
| RMS (angles) (°) | 1.10 |
| Ramachandran favored (%) | 93.98 |
| Ramachandran allowed (%) | 5.59 |
| Ramachandran outliers (%) | 0.43 |
| Rotamer outliers (%) | 0.00 |
| Clashscore | 6.39 |
| Average B-factor ($Å^2$) | 47.85 |
| macromolecules | 47.85 |
| ligands | 58.92 |
| Number of TLS groups | 1 |

**Table S2.** Primers used for constructing the yMGL mutants.

| Primers | Sequences (5´-3´) |
|---|---|
| yMGL-Y61A-F | ATACGTTgcgAGCCGTATTGCGCACCCGACCA |
| yMGL-Y61A -R | TACGGCTcgcAACGTATTCGTTGGTGCCATACG |
| yMGL-R63A -F | TTATAGCgcgATTGCGCACCCGACCACCAGCC |
| yMGL-R63A -R | GCGCAATcgcGCTATAAACGTATTCGTTGGTGCC |
| yMGL-Y88A -F | TGGTTgcgAACAACGGTCTGGCGGCGTTCACC |
| yMGL-Y88A -R | ACCGTTGTTcgcAACCACCGGAAATTTGTTGTTG |
| yMGL-F95A -F | CGgcgACCGCGCTGATCGTGCACGTTAACCCGAA |
| yMGL-F95A -R | ACGATCAGCGCGGTcgcCGCCGCCAGACCGTTGTT |
| yMGL-Y113A -F | TTATTGGCGAAGCGgcgCACGGTTGCCACGCTGTG |
| yMGL-Y113A-R | GTGcgcCGCTTCGCCAATAAACAGACGCTTCGGGT |
| yMGL-A118S -F | CACagcGTGAGCGACATCTTTAAACGTATTCG |
| yMGL-A118S -R | AAGATGTCGCTCACgctGTGGCAACCGTGGTACGC |
| yMGL-V119A-F | CTgcgAGCGACATCTTTAAACGTATTCGTTAT |
| yMGL-V119A -R | AAAGATGTCGCTcgcAGCGTGGCAACCGTGGTA |
| yMGL-D179A-F | ATTCTGAGCGTGgcgGCGACCTTTGCGCCGCCG |
| yMGL-D179A-R | GCcgcCACGCTCAGAATGCCACCTTTCTTGTG |
| yMGL-T181A-F | TGGATGCGgcgTTTGCGCCGCCGCCGCTGCAG |
| yMGL-T181A-R | CGCAAAcgcCGCATCCACGCTCAGAATGCCAC |
| yMGL-S201A-F | ACATCGTTATGCACgcgGCGACCAAGTATTTTGGTGGC |
| yMGL-S201A-R | CGCcgcGTGCATAACGATGTCCGCGCCAAAATCGA |

| | |
|---|---|
| yMGL-K204A-F | CGACCgcgTATTTTGGTGGCCACAGCGATCTG |
| yMGL-K204A-R | ACCAAAATAcgcGGTCGCGCTGTGCATAACGA |
| yMGL-L213A-F | ATCTGgcgGCGGGCATTCTGGTGGTTAAAACC |
| yMGL-L213A-R | AATGCCCGCcgcCAGATCGCTGTGGCCACCAA |
| yMGL-R232A-F | TTGGTGATgcgAGCTTCCTGGGTAGCGGCCCG |
| yMGL-R232A-R | GAAGCTcgcATCACCAACCAGTTGGTCCGCTT |
| yMGL-L235A-F | TCGTAGCTTCgcgGGTAGCGGCCCGGGTAACC |
| yMGL-L235A-R | TACCcgcGAAGCTACGATCACCAACCAGTTGG |
| yMGL-G236A-F | TCGTAGCTTCCTGgcgAGCGGCCCGGGTAACCTG |
| yMGL-G236A-R | CTcgcCAGGAAGCTACGATCACCAACCAGTTGG |
| yMGL-L242A-F | GGGTAACgcgGAAAGCTGGCTGCTGCTGCGTA |
| yMGL-L242A-R | AGCTTTCcgcGTTACCCGGGCCGCTACCCAGG |
| yMGL-E243A-F | TAACCTGgcgAGCTGGCTGCTGCTGCGTAGCC |
| yMGL-E243A-R | GCCAGCTcgcCAGGTTACCCGGGCCGCTACCC |
| yMGL-S244A-F | AACCTGGAAgcgTGGCTGCTGCTGCGTAGCCT |
| yMGL-S244A-R | AGCCAcgcTTCCAGGTTACCCGGGCCGCTACC |
| yMGL-S333A-F | TTAACCACGCGACCgcgCTGGGTGGCGTTGAAAGCC |
| yMGL-S333A-R | AGcgcGGTCGCGTGGTTAAAGAATTTCAGCTTGC |
| yMGL-S347A-F | TCTGATGgcgGACGCGACCACCAACCCGGCGT |
| yMGL-S347A-R | TCGCGTCcgcCATCAGACGCCACTCGATCAGG |
| yMGL-R357A-F | TATCTGgcgGTGAGCGTTGGCGTGGAAGACGC |
| yMGL-R357A-R | ACGCTCACcgcCAGATACGCCGGGTTGGTGGT |
| hMGL (F58Y/E339V/S340A)-F | TTgcgCTGGGTGGTTTTGAAAGTCTGGCGGAA |

| | |
|---|---|
| hMGL (F58Y/E339V/S340A)-R | AAAACCACCCAGcgcAACTGCCAGGGTAAACAGTTTCA |
| CTD[1]-dw-F | CGAGATCAAGAGGACGCTTGTC |
| CTD[1]-dw-R | CTCCGATCTCTTCGACACACTATTGACTCTGTATCACGTGC |
| CTD[2] -up-F | TGATACAGAGTCAATAGTGTGTCGAAGAGATCGGAGCAGTA |
| CTD[2]-dw-R | GTCACACCACTACCATGACCTC |
| P1 | GTCAAGTCTAAGGATCACGTGC |
| P2 | GACCTTCCAACTATACCCCACAC |
